# Vascular dilation modulates brain haematoma expansion in larval zebrafish

**DOI:** 10.64898/2026.03.27.714814

**Authors:** Victor S Tapia, Tom Hardy, Daisy Flatman, Abigail Bennington, Frances Hedley, Poornima Geemon, Catherine B Lawrence, Paul R Kasher

## Abstract

Intracerebral haemorrhage (ICH) is a severe form of stroke with high morbidity and mortality rates. For survivors, acute haematoma expansion strongly determines neurological outcome. Although blood pressure reduction is widely investigated as a strategy to limit haematoma growth, the haemodynamic mechanisms regulating haemorrhage development remain poorly understood. Zebrafish provide a tractable *in vivo* model to study cerebrovascular biology and spontaneous ICH, yet the contribution of vascular regulation to haemorrhage onset and expansion has not been explored in this species. Here, we investigated whether pharmacological modulation of vascular dilation influences ICH development in zebrafish larvae. We first characterised vascular changes during the developmental window in which spontaneous ICH occurs and observed increased heart rate and progressive reductions in arterial diameter between 2 and 3 days post-fertilisation, suggesting increased vascular resistance. We then tested whether vasoconstriction promotes haemorrhage using angiotensin II, which induced systemic and cerebrovascular vasoconstriction but did not increase ICH incidence or haematoma size in two independent ICH models. In contrast, pharmacological vasodilation using sodium nitroprusside or isoproterenol significantly reduced haematoma size in a high-incidence model of atorvastatin-induced ICH. Live imaging of cerebral blood flow revealed that vasodilation was associated with the confinement of red blood cells around affected vessels rather than dispersing into the brain ventricles. Together, these findings indicate that vascular dilation modulates haemorrhage progression in zebrafish ICH and establish this model as a platform to investigate haemodynamic mechanisms regulating haematoma expansion.

## Introduction

Intracerebral haemorrhage (ICH) is the deadliest form of stroke, caused by the spontaneous rupture of cerebral vessels and bleeding into the brain parenchyma^1^. Following the initial rupture, continued bleeding and haematoma expansion frequently occur and represent major determinants of neurological outcome and mortality in ICH patients^1^. Blood pressure plays a central role in both the onset and progression of haematoma expansion. Chronic hypertension is a major risk factor, contributing to hypertensive angiopathy and cerebral small vessel disease (cSVD)^2^, and acute hypertensive episodes are also recognised triggers of spontaneous ICH^3,4^. After ICH onset, acute blood pressure reduction remains one of the most actively investigated therapeutic strategies to limit haematoma expansion and improve outcomes^5,6^. Thus, modulation of blood pressure represents a key pharmacological target both for preventing ICH and for limiting haematoma expansion after haemorrhage.

Animal models are essential for understanding how haemodynamic forces influence brain vessel rupture and haematoma expansion. Both chronic and acute hypertensive stimuli have been modelled in rodents. Genetic and pharmacologically induced models have shown that chronic hypertension induces cerebrovascular damage and promotes spontaneous cerebral microbleeds^7-9^. In hypertensive animals, a secondary acute hypertensive stimulus, such as angiotensin II (Ang II) or norepinephrine, can further trigger spontaneous brain microbleeds^10^. Regulation of haematoma expansion by blood pressure has also been investigated using the collagenase-induced ICH model in rats. In this model, hypertensive animals exhibit increased haematoma size compared to normotensive controls^11,12^, while pharmacological reduction of blood pressure reduces haemorrhage volume and improves behavioural outcomes in hypertensive rats^13^.

Zebrafish provide a complementary model to rodents for translational ICH research, offering a high-throughput system for genetic and pharmacological studies^14^. We and others have shown that zebrafish are a valuable system to study genetic risk factors contributing to spontaneous ICH, including monogenic cerebral small vessel disease (cSVD) and interferonopathies^15-19^, as well as mechanisms of neuroprotection following haemorrhage^20-22^. Notably, cerebrovascular dilation has been observed prior to pharmacologically induced ICH in zebrafish^23^, and increased cerebrovascular diameter has also been reported in a zebrafish *col4a1* crispant model of cSVD, which also shows spontaneous ICH^15^. This suggests that vascular weakness and dilation may precede brain vessel rupture. The optical accessibility and rapid pharmacological manipulation possible in zebrafish larvae provide a unique opportunity to investigate how haemodynamic changes influence haemorrhage development *in vivo*. However, whether haemodynamic forces and vascular resistance influence vessel rupture or regulate haematoma expansion after ICH onset in zebrafish larvae remains unknown.

Here, we investigated whether pharmacological modulation of vascular dilation influences ICH development in zebrafish larvae. We tested whether vasoconstriction increases ICH incidence or haematoma expansion in models with low spontaneous haemorrhage rates, and whether vasodilation limits haematoma growth in a model with high spontaneous ICH incidence. While AngII-induced vasoconstriction did not affect the onset or expansion of haemorrhage, pharmacological vasodilation using sodium nitroprusside (SNP) and isoproterenol (Iso) significantly reduced haematoma size. These findings indicate that haemodynamic modulation influences the progression of ICH in zebrafish, with vasodilation limiting blood extravasation. Together, these results establish zebrafish as a tractable *in vivo* model to investigate haemodynamic mechanisms regulating ICH.

## Results

### Vascular changes during ICH onset

Spontaneous ICH models in zebrafish have been described with an onset between 1.5 and 3 days post-fertilisation (dpf). Pharmacological inhibition of 3-hydroxy-3-methylglutaryl-coenzyme A reductase (hmgcr) using atorvastatin (ATV) induces ICH with an onset between 1.5 and 2 dpf^23,29^. In addition, the cerebral small vessel disease (cSVD) *col4a1* crispant model develops spontaneous ICH between 2 and 3 dpf^15^. Given that these models develop haemorrhage within a similar developmental window, we first characterised vascular changes during this critical period (2–4 dpf; Fig. 1A) in WT larvae before assessing the effects of pharmacological modulation of vascular dilation. Using the endothelial reporter line *Tg(kdrl:eGFP)*, we measured heart rate and dorsal aorta (DA) diameter as indicators of systemic vascular changes and analysed cerebrovascular tone using the basilar artery (BA) diameter as a representative vessel (Fig. 1B). Heart rate increased significantly between 2 and 3 dpf (22% increase on average), and remained stable at 4 dpf (Fig. 1C,D). In contrast, the diameter of the DA progressively decreased during this period, with a 23% reduction from 2 to 3 dpf and a further 19% decrease from 3 to 4 dpf (Fig. 1E,F). Within the brain vasculature, we focused on the BA, which represents the principal artery supplying the hindbrain^30^, because it maintains a consistent dorsal longitudinal position during development (Fig. 1G,H). The BA diameter decreased, albeit non-significantly, by approximately 10% between 2 and 3 dpf and remained stable thereafter (Fig. 1H,I). Together, these data indicate that the developmental window (2-3 dpf) during which ATV-induced and *col4a1*-associated spontaneous ICH occurs coincides with an increase in heart rate and a reduction in arterial diameter, suggesting that ICH onset is temporally associated with increased vascular resistance during early larval development.

**Figure 1.**
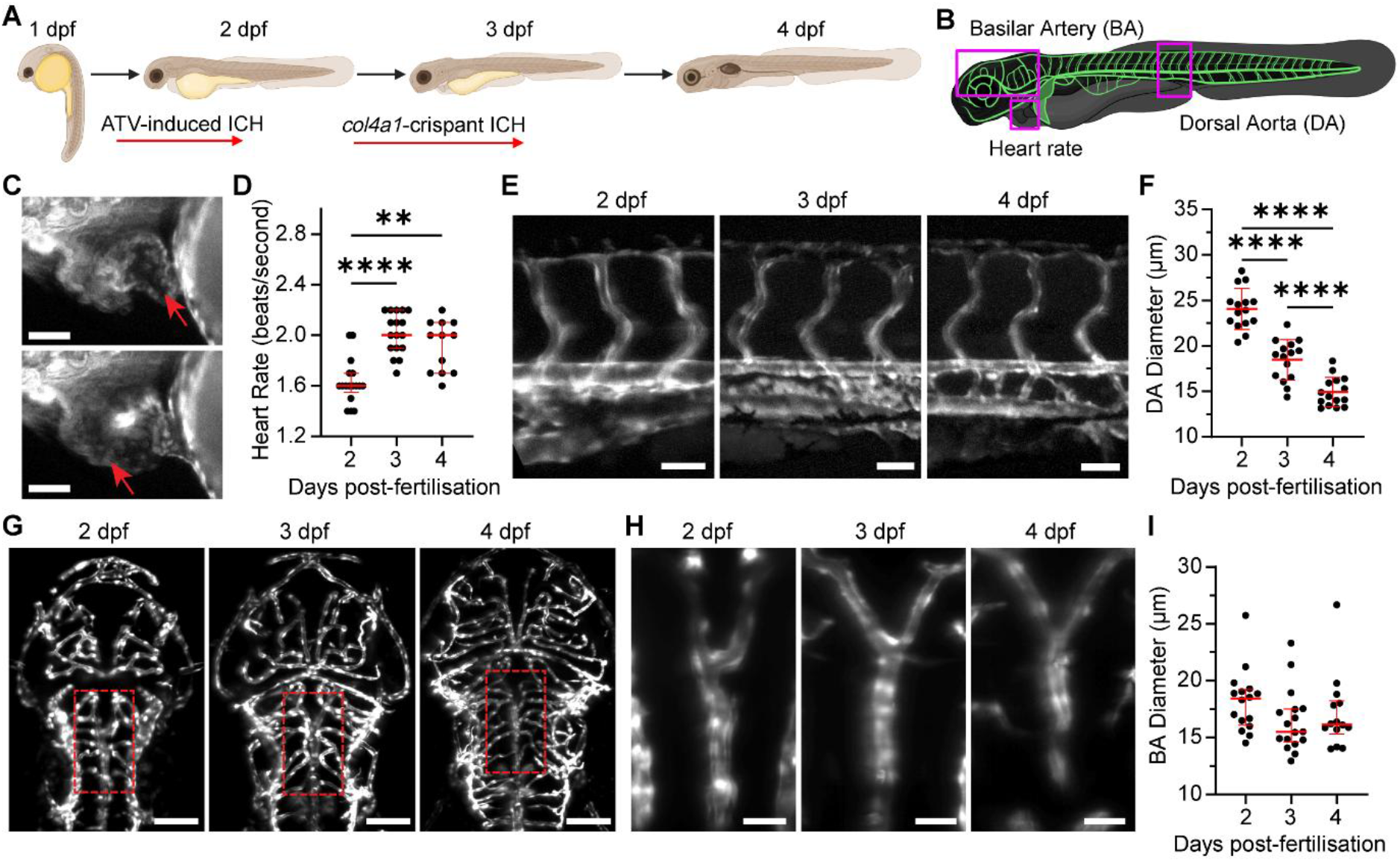
Cardiovascular changes across early larval development. (A) Schematic showing the onset (red arrows) of intracerebral haemorrhage (ICH) in atorvastatin (ATV) and *col4a1* crispant models during early zebrafish development. (B) Schematic of vascular parameters analysed in *Tg(kdrl:EGFP)* larvae across 2-4 dpf. (C) Representative images of the heart at 3 dpf showing the relaxed atrium (top, red arrow) and relaxed ventricle (bottom, red arrow). (D) Quantification of heart rate 2-4 dpf. (E–F) Representative images (E) and quantification (F) of dorsal aorta (DA) diameter at 2-4 dpf. (G) Representative images of cerebral vasculature at 2-4 dpf. Dashed red boxes indicate the position of the basilar artery (BA) shown in (H). (H–I) Representative images (H) and quantification (I) of BA diameter at 2–4 dpf. Scale bars: 50 μm (C, E, G) and 25 μm (H). Each dot represents one larva. Statistics were performed using the Kruskal– Wallis test with Dunn’s post hoc multiple comparison (D, I) or one-way ANOVA with Holm–Šídák’s post hoc multiple comparison (F). Data are presented as median ± IQR (D, I) or mean ± SD (F).**P < 0.01, ****P < 0.0001

### AngII induces vasoconstriction with no effect on brain haematoma dynamics

Next, we tested whether pharmacologically induced vasoconstriction would increase ICH incidence or promote haematoma expansion. Immersion in AngII has previously been shown to induce DA vasoconstriction, in zebrafish larvae at 7-8 dpf (1 μM)^31^, and modulate renal function at 4-5 dpf (1-5 μM)^32,33^. Based on these reported concentrations, we initially tested AngII at 1 μM in 2 dpf embryos and assessed heart rate and DA diameter. However, no vascular effects were observed (data not shown). We therefore evaluated the previously described haemodynamic parameters (Fig. 1) in 2 dpf zebrafish following immersion in AngII at a higher concentration (10 μM) for 3–5 hours. AngII treatment induced a significant 10% increase in heart rate (Fig. 2A), together with significant vasoconstriction of the DA (7% decrease in diameter; Fig. 2B) but no significant changes in the cerebrovascular constriction (BA, Fig. 2C). We also tested a second vasoconstrictor, the thromboxane A_2_ analogue U46619 (U46)^34^. Treatment with U46 (1 or 10 μM) significantly reduced DA diameter in 2 dpf embryos after a 3-hour incubation (Fig. S1A). However, it also reduced heart rate (Fig. S1B), and longer exposure resulted in RBC accumulation in the yolk (Fig. S1C), suggesting impaired systemic circulation. Prolonged exposure also resulted in diminished yolk extension (Fig. S1D). Due to these confounding effects on blood flow, U46 was not further evaluated in the ICH assays.

**Figure 2.**
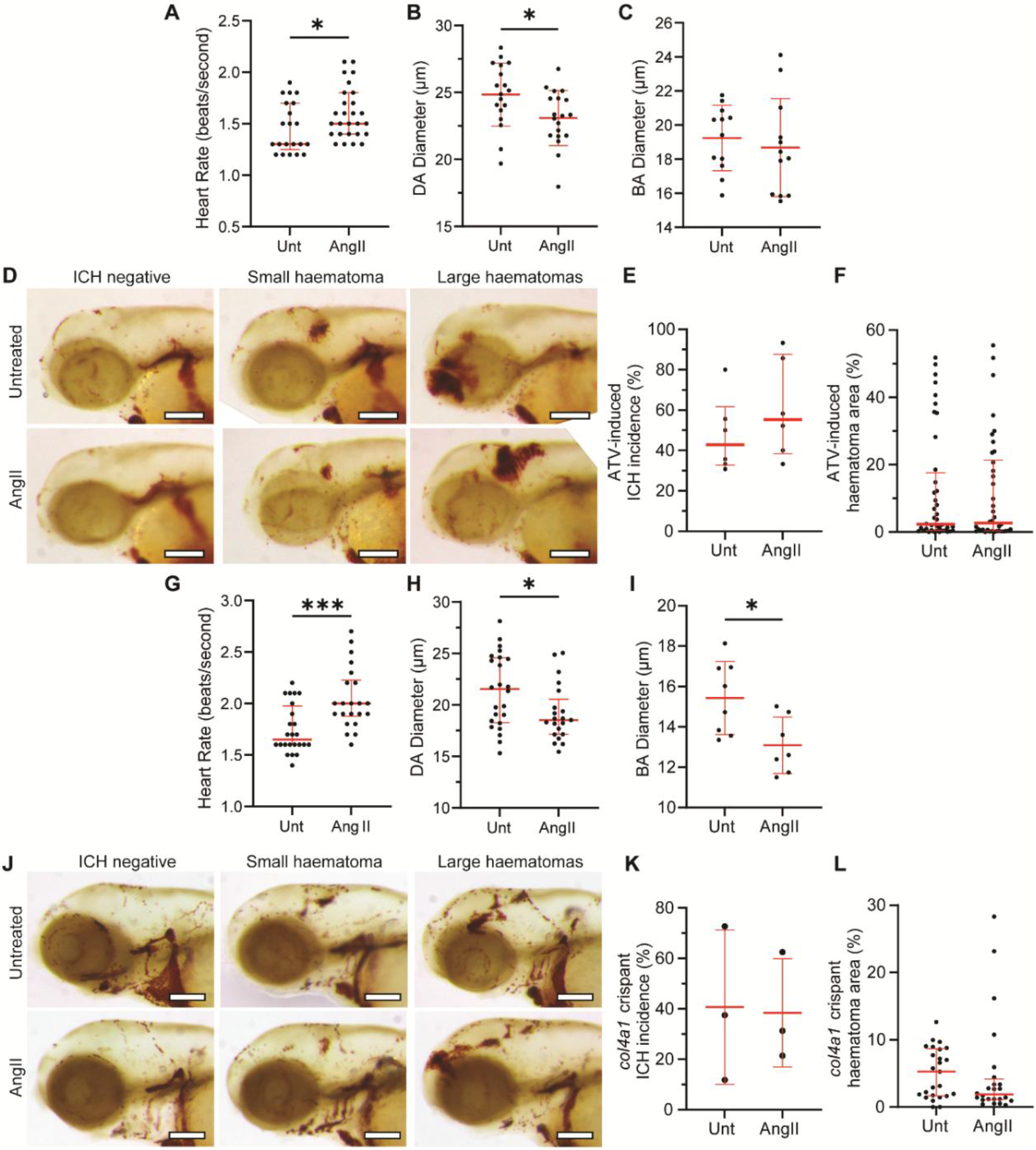
AngII induces vasoconstriction with no effect on ICH. (A–C) At 2 dpf, zebrafish larvae were immersed in angiotensin II (AngII, 10 μM) for 3–5 h. Dorsal aorta (DA) diameter (A), heart rate (B), and basilar artery (BA) diameter (C) were quantified. (D–F) Larvae were immersed in atorvastatin (ATV, 0.5 μM) at 28 hpf, treated with AngII at 32 hpf, and stained with o-dianisidine at 48 hpf. Representative images of intracerebral haemorrhage (ICH) negative larvae and larvae displaying small or large/multiple haematomas are shown (D). ICH incidence (E) and brain haematoma area (F) were quantified. (G–I) At 3 dpf, zebrafish larvae were treated as in panels A– C. DA diameter (G), heart rate (H), and BA diameter (I) were quantified. (J–L) *col4a1* crispants were generated as previously described^15^. Larvae were treated with AngII at 2 dpf and stained with o-dianisidine at 3 dpf. Representative images of ICH negative larvae and larvae displaying small or large/multiple haematomas are shown (J). ICH incidence (K) and brain haematoma area (L) were quantified. Scale bars = 150 μm. Each dot represents one larva (A–C, F–I, L) or one clutch of larvae (E, K). Statistics were performed using the Mann–Whitney test (A, F–H, L), unpaired *t* test (B, C, I), or paired *t* test (E, K). Data are presented as median ± IQR (A, F–H, L) or mean ± SD (B, C, E, I, K). *P* < 0.05, \*\**P* < 0.001.

We next examined whether AngII-mediated vasoconstriction influenced haemorrhage development using a low-incidence model of ATV-induced ICH. Depending on the dose used, ATV concentrations ranging from 0.25 to 1.5 μM induce low to high rates of haemorrhage, corresponding to approximately 25–100% of embryos displaying ICH^16,23^. Zebrafish WT embryos were treated with a low dosis of ATV (0.5 μM) at 28 hours post-fertilisation (hpf), followed by AngII treatment (10 μM) at 32 hpf, and ICH was assessed at 48 hpf using the o-dianisidine assay^28^. ATV treatment induced an average ICH incidence of 47% (percentage of ICH-positive embryos per clutch), and sizes ranging from small to large haematomas (Fig. 2D,E). Although AngII treatment increased the average ICH incidence by 13%, this difference was not statistically significant (Fig. 2E). Similarly, quantification of brain bleed area revealed no change in haematoma size following AngII treatment (Fig. 2F).

To determine whether these findings were consistent across models, we next examined the effects of AngII in a second spontaneous ICH model using *col4a1* crispant embryos, which develop ICH with low incidence and small haematomas at 3 dpf^15^. First, we confirmed that AngII induced vasoconstriction in this developmental stage. Treatment of 3 dpf larvae with AngII (10 μM) for 3–5 hours significantly increased heart rate by 17% (Fig. 2G) and reduced vessel diameter in both the DA (10%) and BA (15%) (Fig. 2H,I). We then generated *col4a1* crispant embryos by injecting a cocktail of 4 gRNAs, as previously described^15^, and treated embryos with AngII at 2 dpf and assessed ICH at 3 dpf. Crispant embryos showed a variable range of ICH incidence and small to large haematomas (Fig. 2J-I), but as in the ATV model, AngII treatment did not increase ICH incidence or haematoma size in this model (Fig. 2J–L). Together, these data show that pharmacological AngII treatment induces systemic and cerebrovascular vasoconstriction in zebrafish larvae but does not modulate ICH development.

### Vasodilators reduce haematoma size in a high-incidence ICH model

Next, we tested whether pharmacologically induced vasodilation would decrease ICH incidence or reduce haematoma expansion. We first evaluated the vasodilator effects of the nitric oxide donor sodium nitroprusside (SNP), which has previously been shown to induce vasodilation in 4-5 dpf zebrafish larvae at 1 μM^35,36^. We treated 2 dpf zebrafish with SNP at 1 or 10 μM for 3-5 hours. Both concentrations significantly increased DA diameter by 7–10% (Fig. 3A), without affecting heart rate (Fig. 3B). We next evaluated isoproterenol (Iso), a β^2^-adrenergic agonist that induces peripheral vasodilation while increasing heart rate^37^. In zebrafish, isoproterenol has been shown to increase heart rate at 10 μM^38^, although its effects on vascular diameter have not previously been characterised. Treatment of 2 dpf larvae with Iso (10 or 100 μM) for 3-5 hours increased DA diameter, although this was only significant at 100 μM, producing a 15% increase (Fig. 3C). As expected, Iso also significantly increased heart rate by approximately 24% at both concentrations tested (Fig. 3D). We further assessed the effects of both vasodilators on the cerebrovasculature using the BA. Neither of them significantly increased BA diameter (Fig. 3E), indicating the vasodilative effects are systemic.

**Figure 3.**
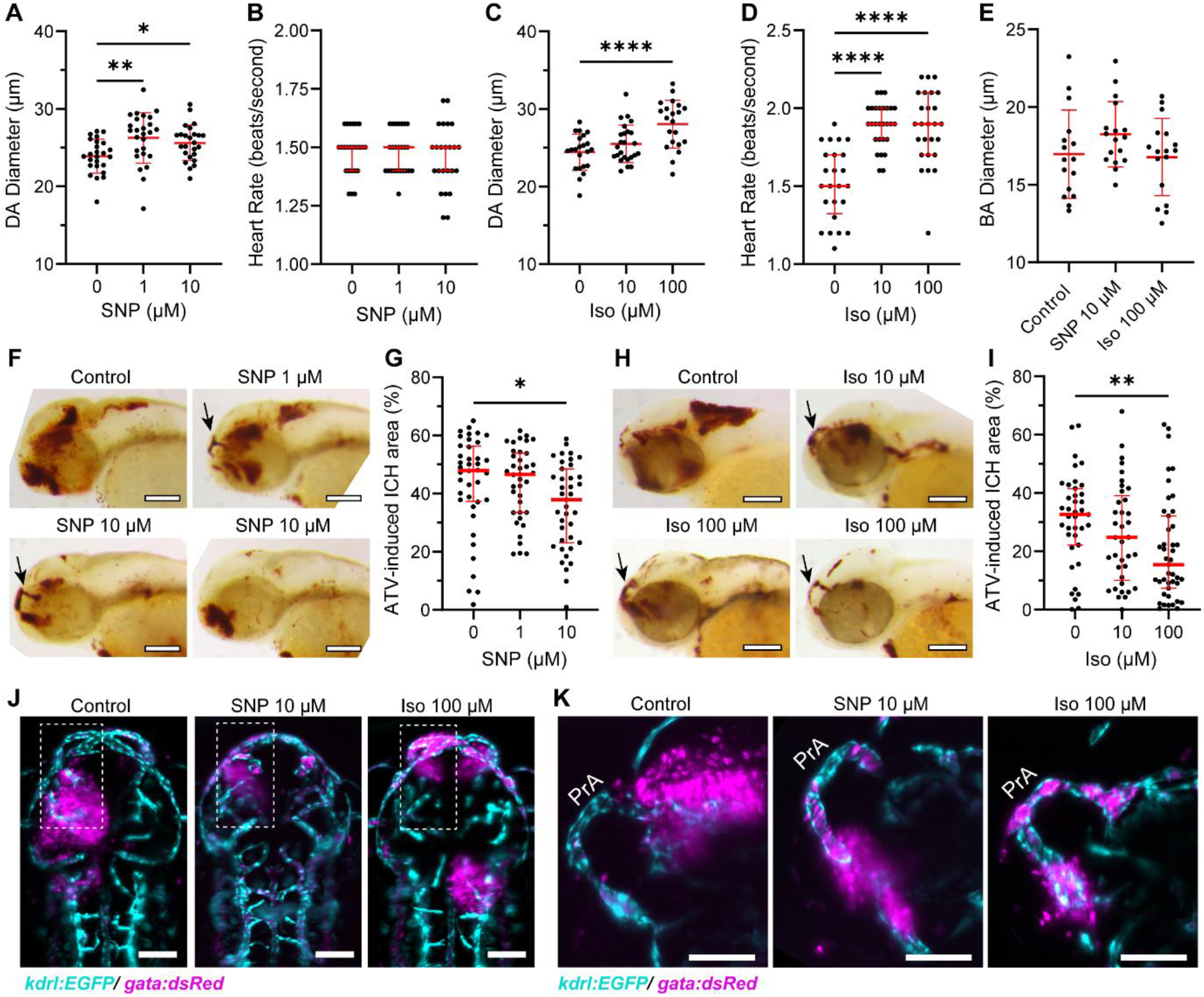
Sodium nitroprusside and isoproterenol induce vasodilation and reduce haematoma size. (A–E) At 2 dpf, WT zebrafish larvae were immersed in sodium nitroprusside (SNP, 1–10 μM) or isoproterenol (Iso, 10–100 μM) for 3–5 h. Dorsal aorta (DA) diameter (A, C), heart rate (B, D), and basilar artery (BA) diameter (E) were quantified. (F–I) WT larvae were immersed in atorvastatin (ATV, 1 μM) at 28 hpf, treated with SNP (F, G) or Iso (H, I) at 32 hpf, and stained with o-dianisidine at 48 hpf. Representative images of control intracerebral haemorrhage (ICH) positive larvae and SNP or Iso treated larvae displaying bulged prosencephalic arteries (PrA) (black arrows) or reduced haematomas are shown (F, H). Brain haematoma area was quantified (G, I). (J–K) *Tg(kdrl:EGFP; gata1:dsRed*) zebrafish larvae were treated as in panels F-I and imaged using light-sheet microscopy. Representative maximum intensity projections of the brain from a dorsal view (J) or of the PrA from a lateral view (K) are shown. Dotted squares in J indicate the region displayed in K. Scale bars = 150 μm (F, H), 75 μm (J, K). Each dot represents one larva. Statistics were performed using one-way ANOVA with Dunnett’s post hoc multiple comparison test versus the control group (A, C, E) or Kruskal-Wallis test with Dunn’s post hoc multiple comparison versus the control group (B, D, G, I). Data are mean ± SD (A, C, E) or median ± IQR (B, D, G, I). *P < 0.05, **P < 0.01, ****P < 0.0001.

We next examined whether these vasodilators influenced haemorrhage development using a high-incidence model of ATV-induced ICH. Zebrafish embryos were treated with ATV (1.0 μM, a higher concentration than used in Fig. 2) at 28 hpf, followed by SNP treatment (10 μM) at 32 hpf, and ICH was assessed at 48 hpf. ATV treatment induced an average ICH incidence of 90% (Fig. S2A), with large and multiple haemorrhages observed in the brain (Fig. 3F). SNP treatment did not reduce ICH incidence (Fig. S2A), but significantly reduced haematoma area at 10 μM (Fig. 3F,G). Notably, this reduction was associated with RBCs localised around the cerebrovasculature rather than accumulating within the brain. This effect was particularly evident in the prosencephalic artery (PrA) (Fig. 3F). A similar effect was observed following Iso treatment. Iso did not reduce ICH incidence (Fig. S2B), but significantly reduced haematoma area at 100 μM (Fig. 3H,I). As with SNP treatment, RBC localisation around the brain vessels was frequently observed in Iso-treated embryos (Fig. 3H).

To further examine RBC localisation, we used the endothelial and erythrocyte reporter line *Tg(kdrl:eGFP; gata1:dsRed)*. Whole-brain imaging confirmed that, following ATV treatment, both SNP and Iso significantly reduced the overall haematoma area (dsRed^+^signal) (Fig. 3J). In ATV-treated control embryos, the PrA was typically devoid of RBC, with blood accumulating within the tectum (Fig. 3K). In contrast, in SNP- and Iso-treated embryos displaying reduced haematoma size, the PrA frequently contained accumulated RBCs, which partially extravasated locally around the vessel (Fig. 3K). Together, these observations suggest that while high-dose ATV induces vessel rupture leading to extended haemorrhage, pharmacological vasodilation may limit haematoma expansion. Under these conditions, RBCs remain partially confined inside or near the affected vessels, such as the PrA, rather than dispersing into the brain parenchyma.

## Discussion

In this study, we investigated whether modulation of vascular dilation influences ICH development in zebrafish larvae. We show that the developmental window during which spontaneous ICH occurs coincides with increased heart rate and reduced arterial diameter, consistent with increased vascular resistance during early larval development. Despite this temporal association, pharmacological vasoconstriction using AngII did not increase ICH incidence or haematoma size in two independent models. In contrast, pharmacological vasodilation using SNP or Iso significantly reduced haematoma size in a high-incidence model. Live imaging further revealed that vasodilation was associated with retention of RBCs near the site of vessel rupture rather than widespread dispersion into the brain. Together, these findings identify vascular tone as a regulator of haemorrhage progression rather than initiation in zebrafish ICH.

The most striking observation was that vasodilation reduced haematoma size without affecting ICH incidence, indicating that vessel rupture still occurs but that bleeding is spatially constrained. In vasodilator-treated embryos, RBCs frequently accumulated around affected vessels, particularly the prosencephalic artery (PrA), whereas in untreated conditions blood dispersed more extensively into the brain parenchyma and ventricles. As the vasodilative effects were observed in the DA but not in the brain, this suggests that systemic vasodilation modifies local haemodynamics in a way that limits the propagation of bleeding. A plausible explanation is that reduced vascular resistance lowers intravascular pressure or alters flow patterns at the rupture site, thereby restricting blood extravasation. This interpretation is consistent with clinical observations that acute blood pressure reduction can limit haematoma expansion^5,6^, although the underlying mechanisms remain incompletely defined.

The zebrafish model provides a unique opportunity to directly test these mechanisms in vivo. Its optical accessibility enables high-resolution analysis of blood flow dynamics, including flow velocity and wall shear stress^36,39-41^. Future work integrating live imaging with quantitative haemodynamic measurements and computational modelling will be essential to determine how systemic vasodilation translates into local changes in shear stress and pressure within the cerebrovasculature. In addition, our data raise the possibility that haemodynamic changes interact with coagulation to limit haemorrhage expansion. The bulged PrA phenotype observed following vasodilator treatment suggests that rupture sites may be stabilised more effectively, potentially through enhanced clot formation or retention. Given that the disruption of coagulation pathways induces spontaneous ICH in zebrafish^42-44^, defining how blood flow and haemostasis interact at sites of vessel injury will be an important next step.

A key feature of zebrafish ICH models is that haemorrhage occurs during early vascular development. In this study, pharmacological modulation of vascular tone was readily observed in the dorsal aorta at 2 dpf, consistent with the onset of vascular smooth muscle cell (vSMC) coverage^45^. In contrast, the basilar artery showed limited responsiveness at 2 dpf, with AngII-induced vasoconstriction becoming evident only at 3 dpf. These findings align with the developmental timeline of the cerebrovasculature, in which pericytes associate with cerebral vessels around 2 dpf and vSMCs appear later at 3-3.5 dpf^46,47^. Functional regulation of cerebrovascular tone has been reported only from approximately 4 dpf onwards^36^. Thus, the cerebral vasculature is structurally and functionally immature during the period in which spontaneous ICH occurs, which is likely to influence both haemorrhage susceptibility and responsiveness to haemodynamic perturbations.

Importantly, multiple zebrafish ICH models converge on a similar developmental window between approximately 1.5 and 3 dpf, including pharmacological, genetic, and morpholino-based approaches^18,19,29,48,49^. In the ATV model, vessel rupture has been linked to the onset of blood flow through newly lumenised vessels^50^, suggesting that the initiation of circulation represents a critical mechanical stress. These observations support the concept that early cerebrovascular development is a period of heightened vulnerability, during which increasing haemodynamic forces act on an incompletely stabilised vascular network. As vascular maturation progresses, susceptibility to spontaneous haemorrhage appears to decrease. Consistent with this, novel zebrafish models targeting pericyte function (*pdgfrb* mutant lines) develop aneurysms and brain haemorrhages at later stages^51,52^, indicating that haemorrhage can also arise in adult zebrafish in contexts of mural cell dysfunction.

Despite inducing systemic and, at later stages, cerebrovascular vasoconstriction, AngII did not increase ICH incidence or haematoma size in either the ATV or *col4a1* crispant models. This contrasts with mammalian studies in which acute hypertensive stimuli can trigger or exacerbate cerebral haemorrhage, particularly in the context of chronic hypertension. The lack of effect observed here likely reflects both the immature state of the cerebrovasculature and the acute nature of the intervention. In mammalian systems, prolonged hypertension is a key driver of vascular remodelling and fragility, which predisposes vessels to rupture. Notably, zebrafish can model aspects of chronic cardiovascular stress, as prolonged AngII exposure induces cardiac hypertrophy in adults^53^. Future studies should therefore examine whether sustained vasoconstriction or hypertension-like conditions influence ICH development, particularly in combination with genetic models with cerebrovascular defects in juvenile or adult zebrafish^15,51,52^.

In summary, our findings demonstrate that vascular dilation is a critical regulator of haemorrhage progression in zebrafish ICH models. While acute vasoconstriction does not promote haemorrhage onset, pharmacological vasodilation limits haematoma expansion and alters the spatial distribution of bleeding within the brain. These results establish zebrafish as a powerful *in vivo* system to study haemodynamic regulation of ICH and provide a framework for identifying therapeutic strategies aimed at limiting haematoma expansion.

## Methods

### Zebrafish husbandry

Zebrafish were raised and maintained at the University of Manchester Biological Services Unit under standard conditions as previously described^24^. Adult zebrafish husbandry was approved by the University of Manchester Animal Welfare and Ethical Review Board. All experiments were performed in accordance with UK Home Office regulations (PPL: PP1968962).

Adults were housed in mixed sex tanks with a recirculating water supply maintained at 28°C under a 14 h/10 h light/dark cycle. Wild-type AB and double transgenic *Tg(kdrl:EGFP)/(gata1a:DsRed)*^*sd2* 25,26^ adult zebrafish lines were used. Fertilised eggs were collected following natural spawning and incubated at 28°C in fresh E3 medium. The embryos were staged according to standard guidelines^27^. After termination of experiments, all embryos were killed before protected status (5 dpf) using a lethal dose of 4% tricaine methanesulfonate (MS222; Sigma-Aldrich) and freezing at −20°C.

Embryos were dechorionated and treated with [Asn1, Val5]-angiotensin II (AngII; Sigma-Aldrich), U46619 (U46; APExBIO), atorvastatin (ATV; Sigma-Aldrich), sodium nitroprusside (SNP; MedchemExpress), or isoproterenol hydrochloride (Iso; Tocris Bioscience). Embryos were maintained at a maximum density of 4 embryos/mL in 12 or 6-well plates (Corning) at 28 °C. Live imaging was performed without anaesthesia at room temperature.

### Heart rate and dorsal aorta imaging

Embryos were positioned laterally in 3% methylcellulose in E3 medium prior to imaging. Fluorescent images were acquired using a Leica M205FA stereofluorescence microscope equipped with a DFC360 FX monochrome digital camera and LAS AF software (version 3.1.0.8587).

Ten-second videos of the embryonic heart were recorded, followed by a single image of the dorsal aorta (DA) centred at the posterior end of the yolk extension. Heart rate was quantified in ImageJ by manual counting to obtain beats per second. DA diameter was measured manually in ImageJ by calculating the average vessel width from five measurements taken between arterioles centred at the posterior end of the yolk extension.

### Cerebrovascular imaging

Embryos were embedded in 1.5% low melt agarose in E3 medium within a glass capillary. Fluorescent images were acquired using a Zeiss Lightsheet Z.1 microscope with Zen software (Zeiss v2.3). A 20x objective with either 0.6x or 1x digital zoom was used to visualise the complete brain vasculature or the prosencephalic artery (PrA), respectively. Z stacks were processed into maximum intensity projections using Zen software and converted to TIFF format. Basilar artery diameter was quantified manually in ImageJ by calculating the average vessel width from multiple measurements along the vessel.

### ICH models

For the ATV induced model, embryos were treated by immersion with ATV (0.5 or 1.0 μM) at 28 hpf and subsequently treated with AngII, SNP, or Iso at specified concentrations at 32 hpf. At 48 hpf, embryos were either stained for ICH or subjected to live imaging.

The *col4a1* crispant model was generated as previously described^15^ targeting the exons 8, 23, 27, and 40 (exon 8 crRNA = 5′-AGACTCACCGGAGGTCCATA-3′; exon 24 crRNA = 5′- GAACCAGGTATAGGTCGGCC-3′; exon 27 crRNA = 5′- GATGGCTTACCTGGTCGCGC −3′;exon 40 crRNA = 5′- GTCCCCTTTAGGCCCCTCCA −3′). These four crRNAs (3.33 μM each) were annealed with tracrRNA (13.33 μM crRNA:tracrRNA duplex) at 95 °C prior to injection. The *col4a1* targeting cocktail was prepared by mixing the crRNA:tracrRNA duplex (8 μM) with EnGen Spy Cas9 protein (4 μM). Freshly fertilised zygotes were injected at the single cell stage into the yolk or cytoplasm with 1 nL of the *col4a1* targeting cocktail. *col4a1* crispants were treated with AngII at 2 dpf and embryos were stained for ICH at 3 dpf.

### O-dianisidine staining

To evaluate haematoma area, embryos were stained at 2 or 3 dpf using an o-Dianisidine (Sigma-Aldrich) protocol, as previously described^23^. Stained embryos were mounted in 100% glycerol and imaged using a Leica M205FA stereo fluorescence microscope, with a DFC425 colour digital camera and processed using LAS AF software (version 3.1.0.8587). Images were analysed as previously described^28^. The brain region was manually selected in ImageJ, and the threshold tool was used identically in all conditions to quantify brain haemorrhage area.

### Statistical analysis

Statistical analyses were conducted using GraphPad Prism v10.6.1 (GraphPad). Shapiro-Wilk normality tests were used to identify parametric and non-parametric datasets. Parametric data are presented as mean ± standard deviation (SD) and non-parametric and ordinal data as median ± interquartile range (IQR). Individual embryos were considered independent datasets. The variable *n* denotes the number of replicates, which corresponds to individual embryos or, in the case of ICH incidence, a clutch of embryos. To compare two independent groups, the unpaired t-test (parametric test) or the Mann-Whitney test (non-parametric test) was chosen. To compare three independent groups, one-way ANOVA (parametric test) or Kruskal-Wallis test (non-parametric) was chosen. ICH incidence was considered matched for embryo clutches, and paired two-tailed t-test (parametric test) or a Friedman test (non-parametric) were used. Post hoc statistical tests are indicated in the corresponding figure legends. Significance was taken at P < 0.05.

## Acknowledgements

We would like to thank the Biological Services Facility at the University of Manchester for expert animal husbandry and the Bioimaging Core Facility at the University of Manchester for their help with imaging.

## Author Contributions

Conceptualization: V.S.T., T.H. and P.R.K; Methodology: V.S.T., T.H. and D.F.; Formal analysis: V.S.T., T.H., D.F., A.B. and F.H.; Investigation: V.S.T., T.H., D.F., A.B., F.H. and P.G.; Data Curation: V.S.T., T.H., A.B. and F.H.; Visualization: V.S.T. and P.R.K.; Supervision: V.S.T. and P.R.K.; Funding acquisition: C.B.L. and P.R.K.; Writing - Original Draft: V.S.T. and P.R.K.; Writing - Editing: All authors.

