## Supplementary Figures for "Vascular dilation modulates brain haematoma expansion in larval zebrafish"

### Supplementary Information

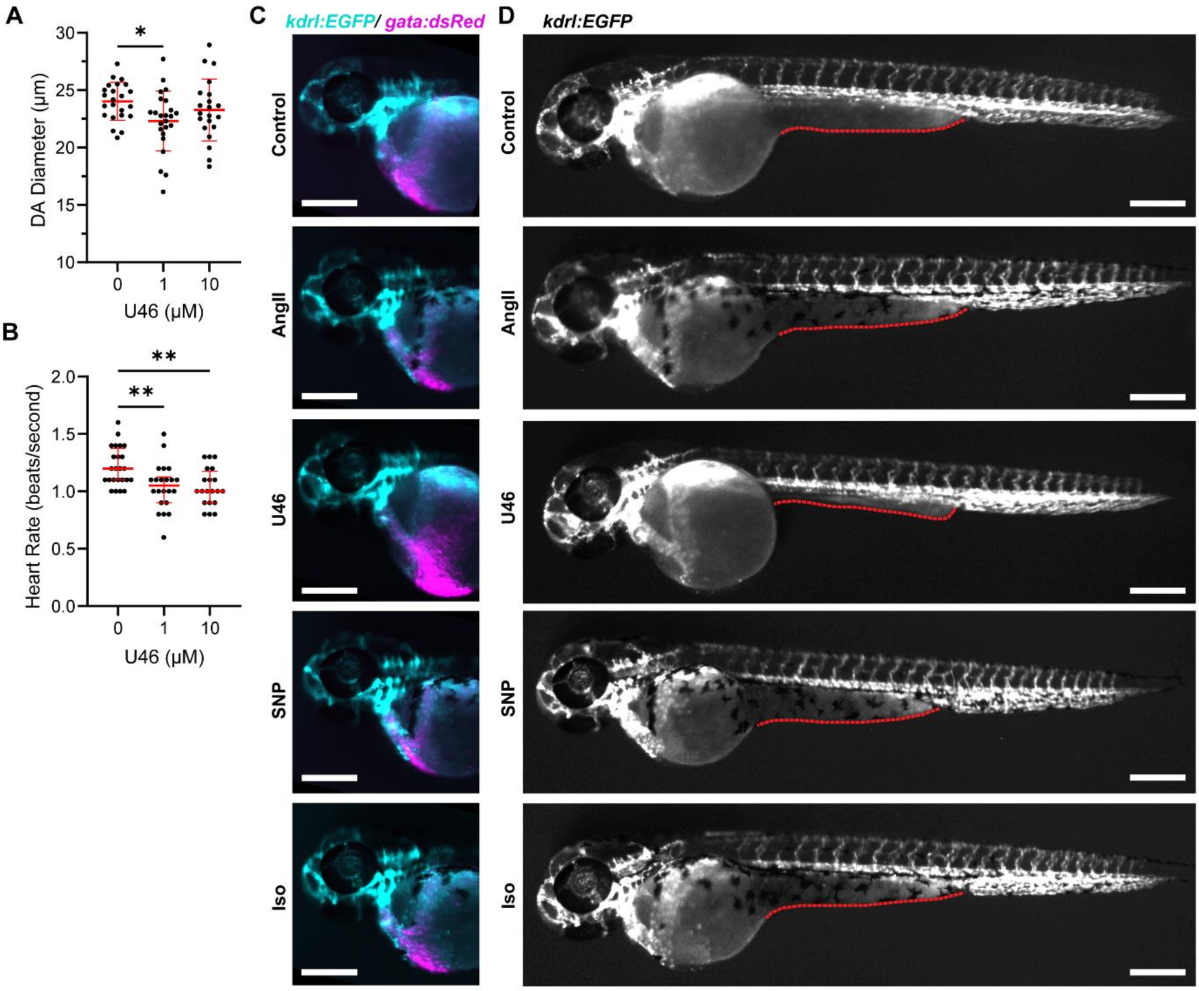


**Supplementary Figure S1. U46619 induces vasoconstriction and developmental defects.** (A–B) At 2 dpf, WT zebrafish larvae were immersed in U46619 (U46, 1–10 µM) for 3–5 h. Dorsal aorta (DA) diameter (A) and heart rate (B) were quantified. (C–D) *Tg(kdrl:EGFP; gata1:dsRed)* larvae were immersed in AngII (10 µM), U46 (1 µM), SNP (10 µM), or Iso (100 µM) at 32 hpf and live-imaged at 50 hpf. Representative images of RBC content in the yolk (C) and whole body (D) are shown. Dotted lines indicate the shape of the yolk extension in (D). Scale bars: 250 µm. Each dot represents one larva. Statistical analysis was performed using one-way ANOVA with Dunnett's post hoc multiple comparison test versus the control group (A), or the Kruskal–Wallis test with Dunn's post hoc multiple comparison test versus the control group (B). Data are presented as mean ± SD (A) or median ± IQR (B). $^*P < 0.05$, $^{**}P < 0.01$.

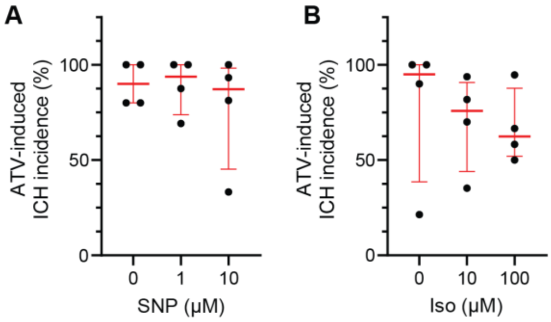


**Supplementary Figure S2. Vasodilators have no effect on ATV-induced ICH incidence.** WT larvae were immersed in atorvastatin (ATV, 1 μM) at 28 hpf, treated with SNP (A) or Iso (B) at 32 hpf, and stained with o-dianisidine at 48 hpf. ICH incidence was quantified. Each dot represents one clutch of larvae. Statistical analysis was performed using the Friedman test with Dunn's post hoc multiple comparison test versus the control group. Data are presented as median ± IQR.
